# NG2 Glia Reprogramming Induces Robust Axonal Regeneration After Spinal Cord Injury

**DOI:** 10.1101/2023.06.14.544792

**Authors:** Wenjiao Tai, Xiaolong Du, Chen Chen, Xiao-Ming Xu, Chun-Li Zhang, Wei Wu

## Abstract

Spinal cord injury (SCI) often leads to neuronal loss, axonal degeneration and behavioral dysfunction. We recently show that in vivo reprogramming of NG2 glia produces new neurons, reduces glial scaring, and ultimately leads to improved function after SCI. By examining endogenous neurons, we here unexpectedly uncover that NG2 glia reprogramming also induces robust axonal regeneration of the corticospinal tract and serotonergic neurons. Such reprogramming-induced axonal regeneration may contribute to the reconstruction of neural networks essential for behavioral recovery.

## INTRODUCTION

Spinal cord injury (SCI) causes the disruption of neural networks, resulting in significant motor and sensory impairments below the affected area. Unfortunately, the adult spinal cord lacks the ability to naturally generate new neurons, and the inability to regenerate damaged axons further hampers neural repair after injury. Our recent study showed the potential of in vivo cell fate reprogramming as a strategy to produce new neurons in the adult mouse spinal cord ^1-3^. However, the impact of this reprogramming process on axonal regeneration of endogenous neurons remains unknown.

Failure of axonal regeneration in the adult spinal cord can be attributed to the reduced intrinsic growth ability of neurons as well as the injury-caused inhibitory microenvironment ^4^. Enhancing neuronal growth capacity, such as by regulating the PI3K/mTOR signaling pathways, shows promise in promoting axonal regeneration after SCI ^5,6^. Similarly, modulating the extrinsic inhibitory microenvironment, such as by degrading the inhibitory chondroitin sulphate proteoglycans (CSPGs) in the lesion scar, also facilitates the regeneration of injured axons across the injury site ^7,8^.

A key cellular component of the injury microenvironment is NG2 glia, which are also known as oligodendrocyte precursor cells ^9^. They remain largely undifferentiated and play dynamic roles during tissue remodeling and repair after SCI ^9-13^. We recently showed that these NG2 glia exhibit latent neurogenic potential in response to SCI and can be further reprogrammed by ectopic SOX2 to become expandable ASCL1+ neural progenitors ^1^. These induced progenitors then generate excitatory and inhibitory neurons that form synaptic connections with local propriospinal neurons and those located in the brain and dorsal root ganglia. Interestingly, SOX2-mediated reprogramming of NG2 glia also changed the microenvironment, indicated by a significant reduction of glial scarring ^1^. Such an observation prompted us to examine the impact of NG2 glia reprogramming on axonal regeneration surrounding the injury site.

## RESULTS

### NG2 glia reprogramming induces robust regeneration of CST axons after SCI

We focused on the corticospinal tract (CST), which is known to be the essential pathway for voluntary movement as well as the most challenging pathway for regeneration after injury ^14-17^.

We employed the 5^th^ cervical vertebra dorsal hemisection (C5-DH) model of SCI since both the dorsal/main tract and lateral tract of descending CST projections could be bilaterally eliminated. Any axons passing through the lesion site were considered as regenerated. We used the same mouse samples as we previously described ^1^. Lentivirus encoding the GFP control, the p75-2 neurotrophic factor, or the SOX2/p75-2 reprogramming factors was delivered to both sides of the lesion one week post injury. The mice were biweekly assessed for behavior. At the end of 26 weeks post SCI, we injected AAV-mCherry as an anterograde tracer into the sensorimotor cortex to label descending CST axons (Fig. 1A-B, S1A-B). We first examined cross-sections of the spinal cord at the far rostral and caudal level to the lesion site. At the far rostral level medullary pyramids, mCherry+ axons were similarly observed among the three treatment groups (Fig. S2A), suggesting comparable AAV injections. On the other hand, mCherry+ axons were not detected at lumbar spinal cord among all groups (Fig. S2B), supporting a complete transection of the CST in our C5-DH injury model.

**Figure 1.**
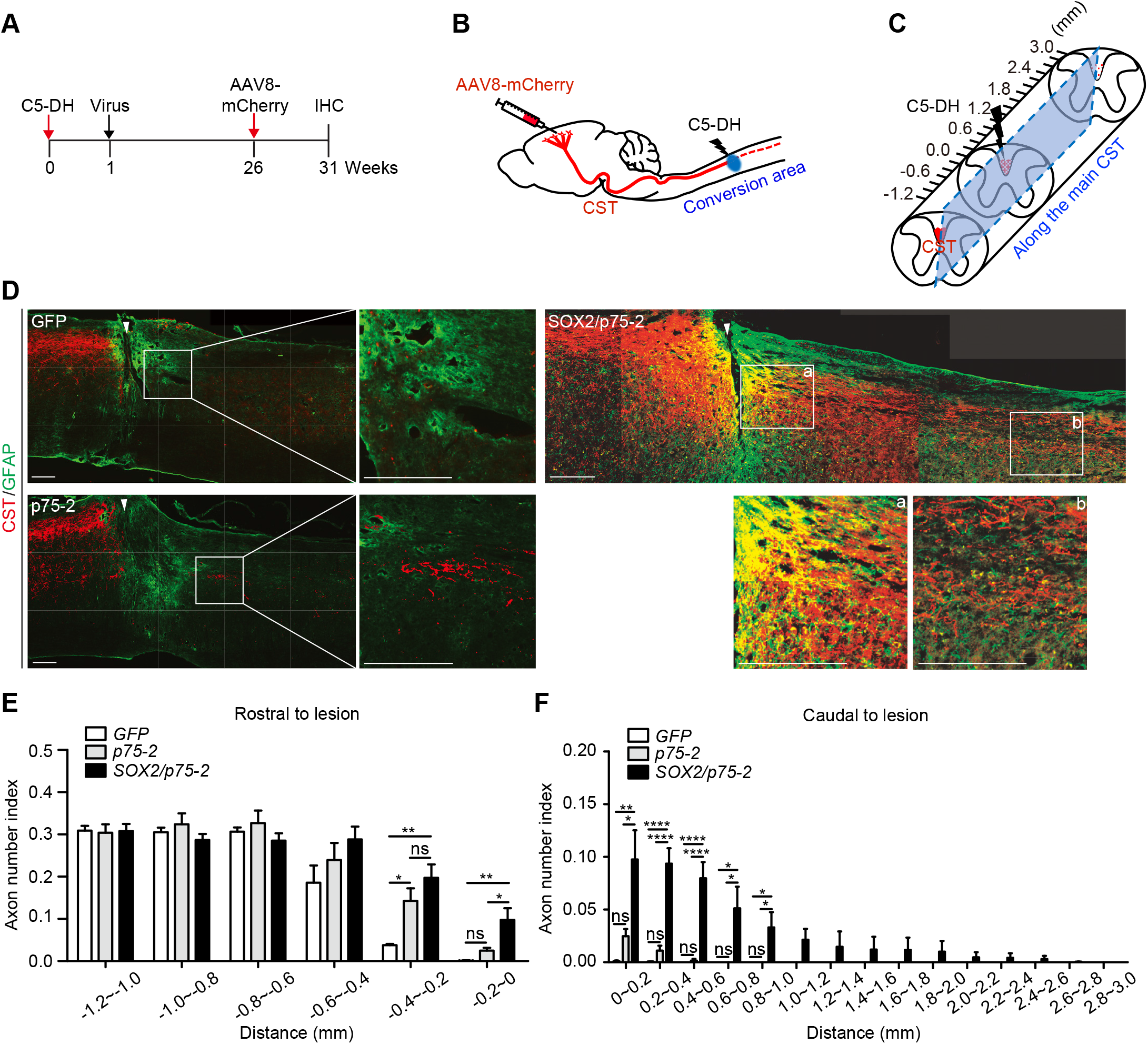
Regeneration of CST axons induced by NG2 glia reprogramming after SCI. (A) Experimental design. (B) Schematic diagram of the injury site and anterograde tracing of the CST axons. (C) Schematic diagram of the spinal cord surrounding the lesion site. The rectangle indicates a sagittal cut through the medial main CST. Axonal densities were measured at both rostral (up to - 1.2 mm) and caudal (up to 3.0 mm) sites to the lesion center (0.0 mm). (D) Confocal images of sagittal sections along the medial main CST. Axons and astrocytes are marked by mCherry and GFAP, respectively. The lesion center is marked by an arrowhead. Enlarged views of the boxed regions are also shown. Scale bars, 200 μm. (E) Quantification of the medial main CST axon density index at sites rostral to the lesion center (*p < 0.05 and **p < 0.01; ns., not significant). (H) Quantification of the medial main CST axon density index at sites caudal to the lesion center (*p < 0.05, **p < 0.01, and ****p < 0.0001; ns., not significant).

We then examined parasagittal sections for mCherry+ axons along both the medial CST sections (Fig. 1C) and those lateral to the main CST (Fig. S1C) across the lesion sites. The GFP group did not show mCherry+ CST axons caudal to the lesion core, consistent with the well-established regeneration failure of these axons in the adult spinal cord (Fig. 1D, S1D). A closer examination around the lesion site rather showed a retraction of mCherry+ axons from the lesion center, a phenomenon known as axon dieback after SCI ^18^. This was quantified as a low axon number index at sites rostral to the lesion center (−0.4 to 0 mm; Fig. 1E, S1E). For the p75-2 group, a few mCherry+ CST axons were detected at the caudal site (Fig. 1D, S1D), though the axon number index was not statistically different from the GFP control (Fig. 1F, S1F). Confirming an axon growth-promoting effect of this neurotrophic factor ^19^, we did find that the axon number index was higher than the GFP control rostral to the lesion center (−0.4 to -0.2 mm; Fig. 1E).

In sharp contrast, the SOX2/p75-2 reprogramming group exhibited very robust regeneration of the mCherry+ main CST axons, extending into regions as far as 2.6 mm caudal to the lesion center (Fig. 1D-F). Extensive growth was also observed for axons lateral to the main CST (Fig. S1D-F). Quantifications confirmed a significantly higher axon number index than either the GFP control or the p75-2 group (Fig. 1E-F, S1F). Together, these results clearly showed that SOX2-mediated NG2 glia reprogramming promotes robust regeneration of CST axons after C5-DH.

### NG2 glia reprogramming promotes regeneration of 5-HT axons after SCI

We also examined the serotonergic pathway for the effect of NG2 glia reprogramming on axonal regeneration. Serotonin (5-HT) plays an important role in modulating the activity of spinal networks involved in locomotor functions. SCI can disrupt the descending serotonergic projections to spinal motor areas, leading to locomotor dysfunction ^20^. In contrast to the CST, the 5-HT axons exhibited increased axonal sprouting rostral to the lesion center in all experimental groups (Fig. 2A), consistent with the other published observations ^21^. When examining the caudal side of the lesion, interestingly, we detected many more 5-HT axons in the SOX2/p75-2 reprogramming group (Fig. 2A). Quantitative analysis revealed a significant increase of the axon intensity index within 2.0 mm caudal to lesion border (Fig. 2B). Such a result indicates that NG2 glia reprogramming promotes regeneration of 5-HT axons after SCI.

**Figure 2.**
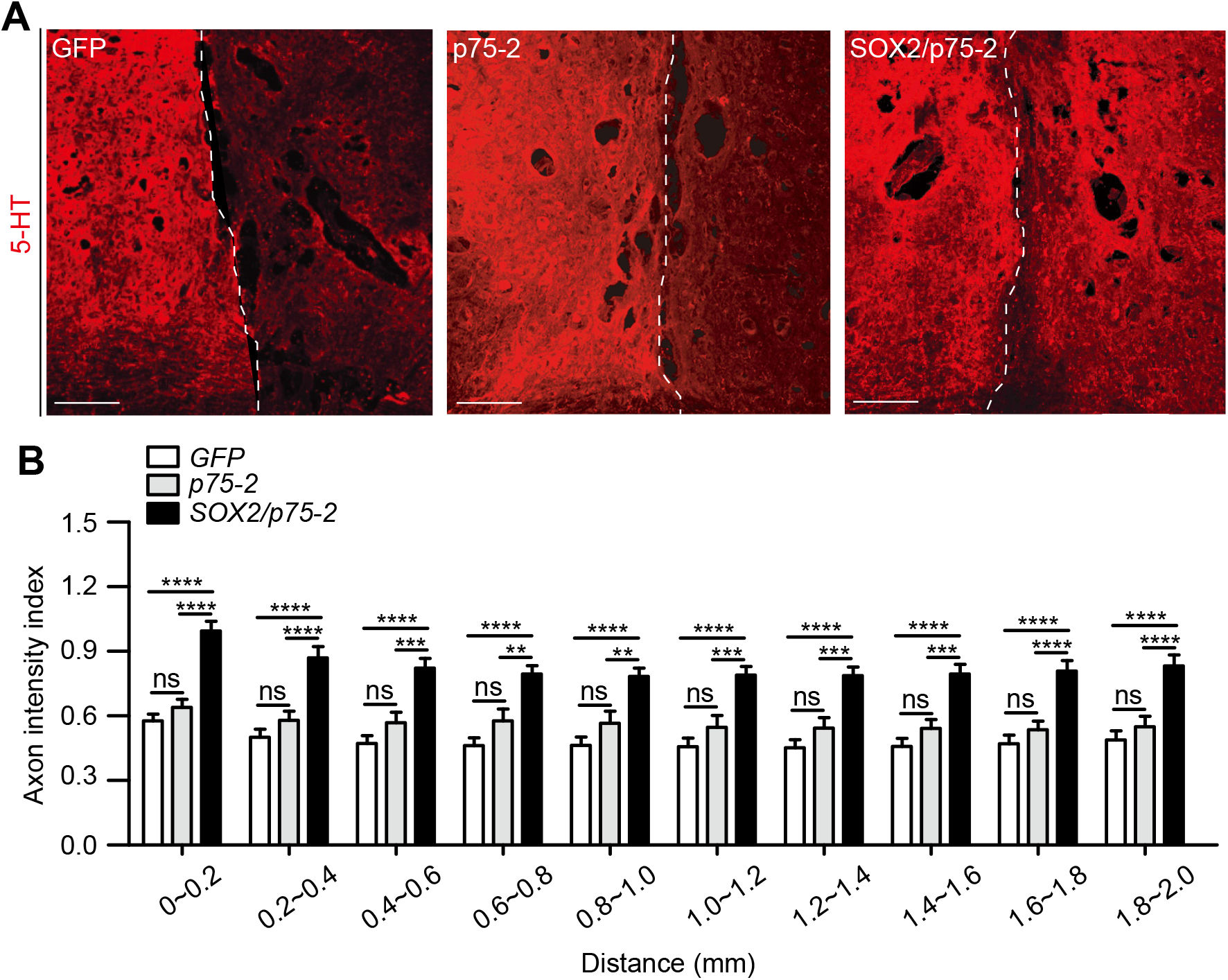
Regeneration of 5-HT axons induced by NG2 glia reprogramming after SCI. (A) Confocal images of 5-HT staining surrounding the lesion center. The dotted line marks the lesion center. Scale bars, 200 μm. (B) Quantification of axonal density index at sites caudal to the lesion center (**p < 0.01, ***p < 0.001, and ****p < 0.0001; ns., not significant).

## DISCUSSION

In vivo reprograming of resident glial cells is emerging as a regenerative strategy for new neurons in the adult central nervous system ^22-27^. For the first time, the results of this study reveal that glia reprogramming also induces robust regeneration of CST axons and 5-HT axons across the lesion after SCI.

Originating from corticospinal neurons in the motor cortex, the CST axons transmit cortical commands to the spinal cord to control movements ^28^. Disruption of CST axons results in severe motor deficits at and below the injury level. 5-HT is a monoamine neurotransmitter synthesized in various populations of brainstem neurons. The 5-HT axons project to all levels of the spinal cord and are important in modulating locomotion. Disruption of the serotonergic pathway produces varying degrees of locomotor dysfunction ^20^. Despite their pivotal roles, adult CST axons and 5-HT axons exhibit limited intrinsic ability to regenerate after SCI ^5,29,30^. How does NG2 glia reprogramming lead to enhanced regeneration of these axons?

Both intrinsic and extrinsic pathways are found to control axonal regeneration ^7,31-34^. For example, modulating the mammalian target of rapamycin (mTOR) pathway in pyramidal neurons induces robust CST regeneration after SCI ^5,28^. Transplantation of neural progenitor cells (NPCs) also enables growth of the CST axons into the grafts and beyond the lesion sites ^35^. This is largely due to that the grafted NPCs revert the injured CST neurons into an immature transcriptional state ^36^. A similar mechanism may be applied to the NG2 reprogramming-initiated axonal regeneration identified in this study, because many ASCL1+ NPCs and DCX+ immature neurons were induced at the injection sites during the reprogramming process ^1^. These NG2 glia-derived NPCs and immature neurons may remodel in a cell non-autonomous manner the transcriptional programs of CST neurons to enable their intrinsic growths.

For decades, the glial scar has been recognized as a critical extrinsic modulator of axonal regeneration. As a major cellular component of the glial scar, NG2 glia rapidly accumulate around the lesion site, leading to increased levels of the NG2 proteoglycan, a key constituent of the inhibitory chondroitin sulfate proteoglycans (CSPGs) that impede axonal regeneration ^9,37-40^. NG2 glia play a key role in remodeling the post-SCI tissue for scar formation and their transient early loss but later repopulation leads to axonal growth into the lesion sites ^13^. Such a role of NG2 glia is supported by our finding of the reduction of scaring tissues after NG2 glia reprogramming ^1^. Of note, NG2 glia reprogramming by SOX2 produces not only new neurons but also an almost equal number of new NG2 glia/oligodendrocytes ^1^. These reprogrammed new glial cells may facilitate axonal regrowth as well as their subsequent remyelination.

Together, the results of this study uncover an unexpected biological function of NG2 glia reprogramming in vivo. Such reprogramming may have the “killing two birds with stone” effect on tissue remodeling and regeneration after SCI. Future studies are warranted to tease out the underlying molecular and cellular mechanisms and further develop the in vivo reprogramming approach as a therapeutic strategy for neural repair.

## EXPERIMENTAL MODEL AND SUBJECT DETAILS

### Animals

We used the same animal samples as described previously ^1^, Briefly, wild-type C57BL/6J mice (JAX: 000664; RRID: IMSR_JAX:000664) were purchased from the Jackson Laboratory. Both adult male and female mice at 2 months of age and older were used for all experiments unless otherwise stated. All mice were housed under a controlled temperature and a 12-h light/dark cycle with free access to water and food in the animal facility. Sample sizes were empirically determined. Animal procedures and protocols were approved by the Institutional Animal Care and Use Committee at UT Southwestern or Indian University School of Medicine.

## METHOD DETAILS

### C5-DH spinal cord injuries model

The C5-DH was conducted following our previous published studies. Briefly, after anesthetization and spine stabilization, a laminectomy was conducted to expose the C5 spinal cord. A transverse durotomy at the interlaminar space was performed using a 30 G needle, which was followed by cutting with a pair of microscissors. The VibraKife, attached to the Louisville Injury System Apparatus (LISA), was used to cut the spinal cord according to our previously published method ^14,16,41^. The blade, vibrating at 1.2 mm wide, was slowly advanced 1.2 mm ventrally from the dorsal surface of the cord, resulting in complete transection of the entire dorsal corticospinal tract CST (dCST) and lateral corticospinal tract CST (lCST). In this injury model (1.2 mm wide, 1.2 mm depth), any labeled CST axons growing beyond the lesion would be interpreted as coming from cut axons via regeneration from the tips or by regenerative sprouting.

### Corticospinal tract tracing

The procedure of tracing the CST axons is similar to our previously published studies ^14,16,17^. Briefly, after exposing the skull, bilateral windows (5 mm in length and 2 mm in width) were created with the medial edges of windows 0.5 mm lateral to the bregma. Using a digital stereotaxic injector (Item: 51709, Stoelting Co. USA), 0.5 μl of AAV8-mCherry (5 × 10^12^ GC/μl) was injected into one of 10 total sites (5 sites/side). The mediolateral (ML) coordination was 1.5 mm lateral to the bregma, the anteroposterior (AP) coordination from the bregma was at -1.0, - 0.25, +0.5, +1.25 and +2 mm; dorsoventral (DV) coordination: 0.5 mm from the cortical surface; rate: 0.1 μl/minute. After each injection was completed, the injector tip was left in place for an additional 5 min to adequate penetration of the tracer. Five weeks later, the mice were anesthetized and perfused with 4% paraformaldehyde for detecting CST distribution in the spinal cord.

### Immunofluorescence staining

Mice were sacrificed and sequentially perfused with ice-cold phosphate-buffered saline (PBS) and 4% (w/v) paraformaldehyde (PFA) in PBS. Tissues were harvested, post-fixed in 4% PFA overnight, dehydrated in 30% sucrose for 2 days. Serial sagittal or cross cryostat-sections were stained with primary antibodies in blocking buffer (3% goat serum or 5% donkey serum in 0.3% PBST) for at least 18 h at 4LJ. The primary antibodies were mouse anti-GFAP (1:2000, Abcam, ab8049) and rabbit anti-serotonin (5-HT, 1:500, Sigma-Aldrich, S5545). Detection was accomplished by incubation with Alexa Fluor-coupled secondary antibodies (Life Technologies/Abcam) diluted in blocking buffer for 2 h at room temperature. The OLYMPUS FLUOVIEW FV1200 confocal microscope was used to visualize the axonal regeneration on the sections (20× and 60×).

### Data analysis

To quantify the fiber number index of CST axons, we followed the established protocol ^6,14,16^ with minor modifications. Briefly, 3-4 sagittal sections from each mouse were analyzed using the ImageJ software. The number of CST axon fragments was counted at defined zones spaced 0.2 mm apart. The axon number was standardized into Axon Number Index, defined as the ratio of the mCherry-labeled CST axon number at a given zone over the total number of axons in the descending CST on a cross section of medullary pyramids. The lesion epicenter on the sagittal section was defined as “0” and all other zones were defined as their distance from the lesion center. To quantify the intensity index of the 5-HT-positive axons rostral and caudal to the lesion site, 3 sagittal sections from each animal were analyzed using ImageJ. In each sagittal section, we drew a (dorsoventral) line starting from the rostral lesion border (0.0 mm), then vertical lines at distance caudal to lesion border was drawn at the distance of 0.2 mm. The axon intensity index was presented as a ratio of axon density at a specific region versus the axon density 1.0 mm rostral to the lesion border. The averaged axon intensity per animal was further processed in GraphPad to generate regional density maps at certain distances from the starting line.

### Quantification and statistical analysis

All data are presented as mean ± SEM. Statistical analyses were carried out using Prism 7 (Graphpad Software). Comparisons between GFP, p75-2 and SOX2/p75-2 groups were performed by One-way ANOVA. A p value < 0.05 was considered significant. Significant differences are indicated by *p < 0.05, **p < 0.01, ***p < 0.001, and ****p < 0.0001.

## ACKNOWLEDGMENTS

We thank members of the Zhang and Wu laboratory for discussions and reagents. C.-L.Z. is a W.W. Caruth, Jr. Scholar in Biomedical Research. The work in the Zhang laboratory was supported by the Decherd Foundation, the Texas Alzheimer’s Research and Care Consortium (TARCC2020), and the NIH (NS099073, NS092616, NS111776, NS117065, and NS088095). The work in the Wu laboratory was supported by the NIH (R01NS103481 and 1R01NS111776) and Indiana Department of Health (ISDH58180).

## AUTHORS’ CONTRIBUTIONS

W.T., W.W., X.-M.X., and C.-L.Z. conceived and designed the experiments. W.T., X.L.D, C.C., and W.W., performed the experiments. W.T., C.-L.Z and W.W. analyzed data. W.T., C.-L.Z. and W.W wrote the manuscript. All authors (except for X.-M.X.) reviewed and approved the manuscript.

## DECLARATION OF INTERESTS

The authors declare no competing interests in the manuscript.

## Notes

### Competing Interest Statement

The authors have declared no competing interest.

