## Supplementary Figures for "NG2 Glia Reprogramming Induces Robust Axonal Regeneration After Spinal Cord Injury"

Sup Figure_1

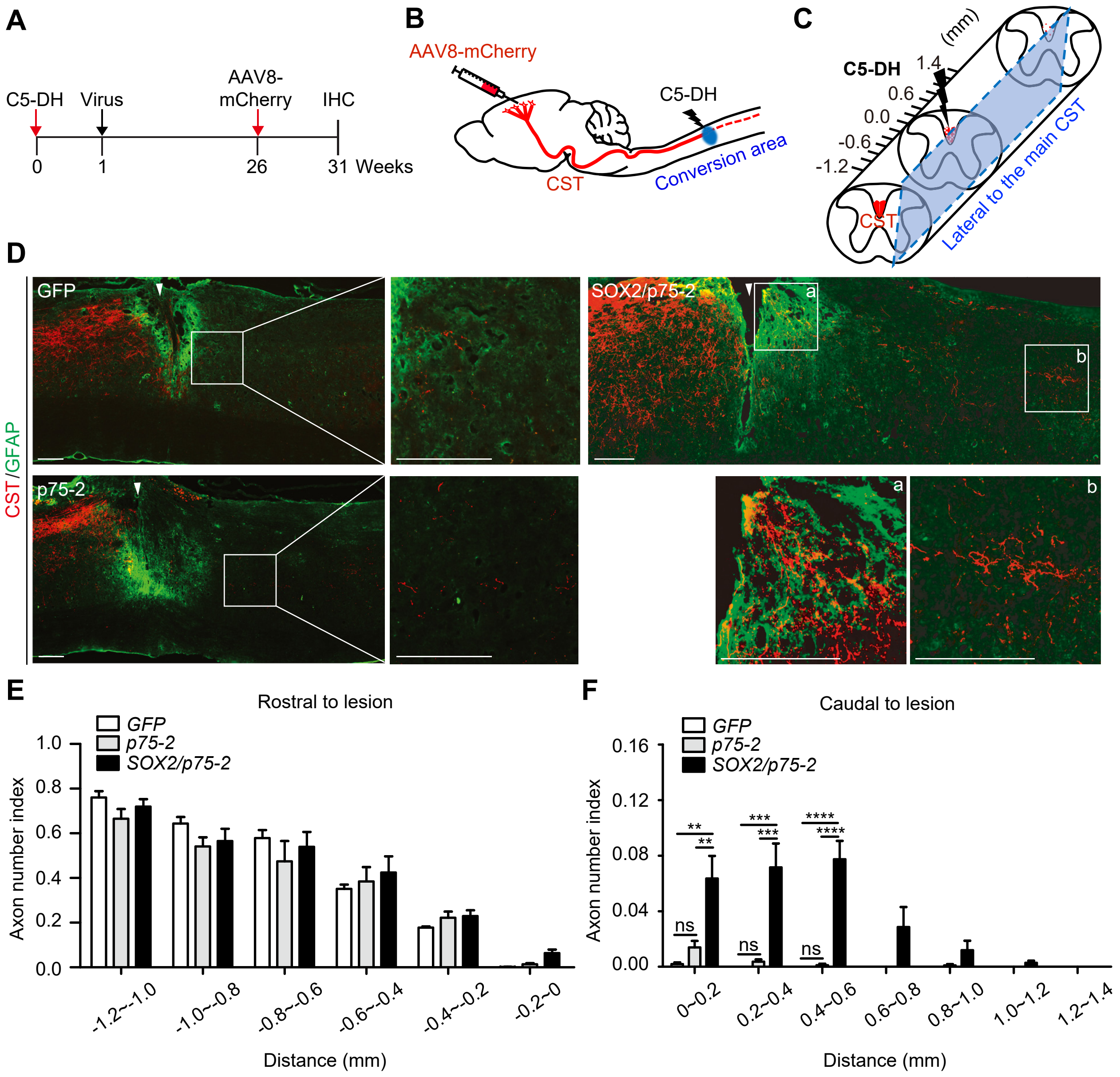


**Figure S1. Regeneration of CST axons induced by NG2 glia reprogramming after SCI.**

(A) Experimental design.

(B) Schematic diagram of the injury site and anterograde tracing of the CST axons.

(C) Schematic diagram of the spinal cord surrounding the lesion site. The rectangle indicates a parasagittal cut through the lateral to the main CST. Axonal densities were measured at both rostral (up to -1.2 mm) and caudal (up to 1.4 mm) sites to the lesion center (0.0 mm).

(D) Confocal images of parasagittal sections along the lateral to main CST. Axons and astrocytes are marked by mCherry and GFAP, respectively. The lesion center is indicated by an arrowhead. Enlarged views of the boxed regions are also shown. Scale bars, 200 µm.

(E) Quantification of the lateral CST axon density index at sites rostral to the lesion center.

(F) Quantification of the lateral CST axon density index at sites caudal to the lesion center (**$p < 0.01$, ***$p < 0.001$, and ****$p < 0.0001$; ns., not significant).

**Sup Figure_2**

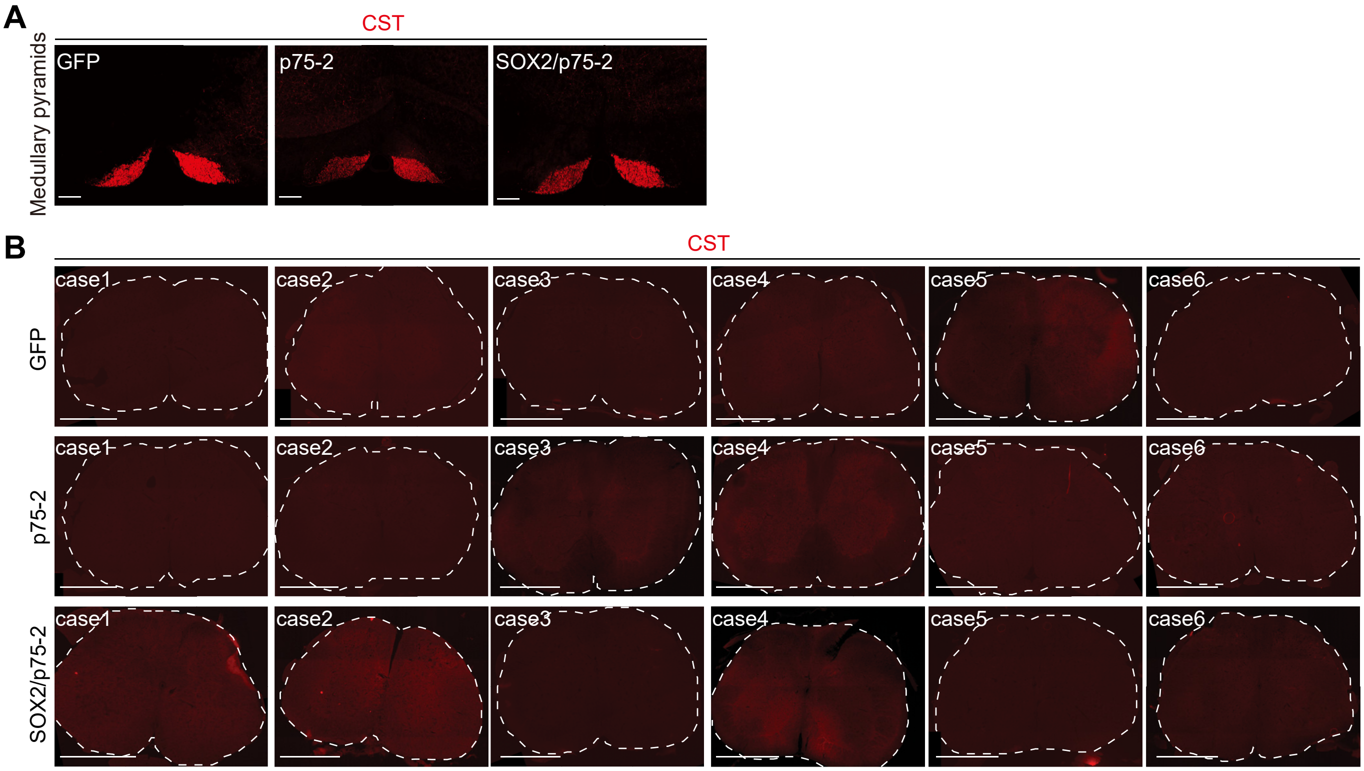


**Figure S2. CST axons at the far rostral and caudal sties to the lesion center.**
(A) Images of transverse sections of traced CST axons at the medullary pyramids. Comparable mCherry+ axons were observed at the far rostral sites. Scale bars, 200 µm.
(B) Images of transverse sections of traced CST axons at the lumbar spinal cord. Each individual animal case is shown. mCherry+ axons were not detected at the far caudal sites. Scale bars, 500 µm.
